# Estimating the number of protein molecules in a plant cell: a quantitative perspective on proteostasis and amino acid homeostasis during progressive drought stress

**DOI:** 10.1101/2020.03.17.995613

**Authors:** Björn Heinemann, Patrick Künzler, Hans-Peter Braun, Tatjana M. Hildebrandt

## Abstract

During dehydration cellular proteostasis as well as amino acid homeostasis are severely challenged, since the decrease in photosynthesis induces massive proteolysis. Thus, we selected progressive drought stress in *Arabidopsis thaliana* as a model to investigate the balance between protein and free amino acid homeostasis on a quantitative level. We analyze the mass protein composition of rosette leaves and estimate, how many protein molecules are present in a plant cell and its subcellular compartments. Under control conditions, an average Arabidopsis mesophyll cell contains about 25 billion protein molecules and 80% of them are localized in the chloroplasts. Severe water deficiency leads to degradation of more than 40% of the leaf proteome and thus causes a drastic shift towards the free amino acid pool. Stress induced proteolysis of half of the 400 million RubisCO hexadecamers present in the chloroplasts of an individual mesophyll cell alone doubles the cellular content in free amino acids. A major fraction of the amino acids released from proteins is channeled into the synthesis of proline as a compatible osmolyte. Complete oxidation of the remaining part as an alternative respiratory substrate can fully compensate the lack of carbohydrates derived from photosynthesis for several hours.

## Introduction

Proteostasis (protein homeostasis) is essential for maintaining normal cellular functions, which rely on an appropriate composition as well as correct folding of the proteome. Plant cells contain several thousand different proteins that are highly diverse not only in terms of their function but also in size and abundance. RubisCO has to be present in large quantities in leaf cells due to its low enzymatic activity and carbon fixation efficiency, whereas hardly detectable amounts of e.g. signaling molecules or transcription factors are sufficient to fulfil their function. The protein composition of other tissues such as roots or seeds again is completely different (Baerenfaller et al. 2008, Mergner et al. 2020). In addition, 1 mg of a large protein such as glutamate synthase contains only 4 nmol active sites compared to 83 nmol for the small protein glutaredoxin. Thus, the investment of resources (energy and nutrients) required for the synthesis of large and/or high abundant proteins is by several magnitudes higher than for small proteins of low abundance.

Not surprisingly, cells contain several sophisticated systems to control proteostasis and recycle the resources needed for new growth. Protein synthesis is catalyzed by the ribosomes in the cytosol, plastids, and mitochondria. The synthesis rate is regulated on different levels in response to the energy status of the cell, e.g. via mRNA availability, the GDP and GTP pools, and posttranslational modification of the ribosome (Merchante et al. 2017). The two major protein recycling systems in eukaryotes are autophagy and the ubiquitin-proteasome system (reviewed by Dikic 2017, Marshall and Vierstra 2018, Vierstra 2009). During autophagy cytoplasmic constituents including large protein and nucleic acid aggregates, lipid bodies, and even entire organelles are sequestered into a double membrane vesicle, the autophagosome, and delivered to the vacuole for breakdown. Thus, autophagy in addition to proteins also digests nucleic acids, lipids, and carbohydrates. Autophagosome formation is controlled by a highly conserved set of 40 autophagy-related (ATG) proteins. They include receptors that recognize specific cellular components and tether them to the enveloping autophagic membrane to target them for destruction. In contrast, the ubiquitin-proteasome system localized in the cytosol catabolizes proteins individually. Substrates are marked for degradation by a poly ubiquitin tag that enables their recognition and hydrolysis by the proteasome, a large protein complex composed of a 20S catalytic core and two regulatory 19S lids. Several molecules of the 8.5 kDa protein ubiquitin are covalently conjugated to a lysine residue of the substrate protein by an enzymatic cascade consisting of ubiquitin activating (E1), conjugating (E2), and ligating (E3) enzymes. Substrate specificity is provided by a high number of different E3 ubiquitin ligases (>1,400 in the Arabidopsis genome). In addition to the bulk degradation systems plants contain hundreds of individual proteases from several unrelated families. They can be grouped into four major classes according to the nature of the nucleophile used for proteolytic cleavage of the peptide bond. Cysteine and serine proteases use a Cys or Ser activated by His as a nucleophile whereas metalloproteases and aspartic proteases activate water using a metal ion or Asp, respectively (van der Hoorn, 2008). Proteases are present in all the different subcellular compartments. Plastids and mitochondria contain distinctive proteolytic systems from prokaryotic origin such as AAA-class, Lon, FtSH and Clp proteases (Nishimura et al. 2016; Kwasniak et al. 2012).

The accumulation of non-functional and misfolded proteins would lead to the formation of large protein aggregates that are detrimental to cellular function (McClellan et al. 2005). Thus, damaged proteins are efficiently detected and eliminated by the two main protein quality control systems, the ubiquitin-proteasome system and autophagy, to avoid proteotoxic stress (Dikic 2017). Even under steady state conditions the turnover rates of individual proteins are highly diverse, a more than 150-fold variation in protein degradation has been reported (Li et al. 2017). The D1 protein localized in the reaction center of photosystem II is replaced on a daily basis since it is frequently damaged by reactive oxygen species as a result of photosynthetic activity. Also, regulatory proteins such as hormone response factors usually have a short half-life to allow rapid responses to a changing environment (Nelson and Millar 2015). In contrast, ribosomal subunits are among the most stable proteins in Arabidopsis and remain functional for several months (Li et al. 2017). Protein stability is defined by different factors such as the physical location of the protein, interactions with cofactors or other proteins, and post-translational modifications (Nelson and Millar 2015).

Proteostasis is closely connected to amino acid homeostasis since protein synthesis requires sufficient supply of loaded t-RNAs whereas proteolysis releases free amino acids. The effect of protein metabolism on the relative contents of free amino acids can be substantial in particular for low abundant amino acids such as the sulfur containing, aromatic, and branched chain amino acids (Hildebrandt 2018). In yeast and animal cells proteasome inhibition leads to cell death, which is primarily caused not by the accumulation of misfolded proteins but by a detrimental deficiency in free amino acids (Suraweera et al. 2012). Apart from serving as building blocks for proteins free amino acids have several additional functions in plant metabolism. They are precursors for the synthesis of secondary metabolites, hormones and signaling molecules, and also act as transport and storage forms for organic nitrogen (Alcázar et al. 2006; Lam et al. 2003; Tzin and Galili 2010). During drought and salt stress Pro and the non-protein amino acid γ-aminobutyric acid (GABA) function as compatible osmolytes (Krasensky and Jonak 2012). Proteolysis is increased in response to adverse environmental conditions to provide amino acids as precursors for these defense related metabolites and also as alternative substrates for ATP production when photosynthesis rates are low (Araujo et al. 2011; Hildebrandt et al. 2015). In the present study we use progressive drought stress in Arabidopsis as a model to investigate the balance between protein and free amino acid homeostasis on a quantitative level. We estimate the molecular as well as the mass protein composition of an average rosette leaf and an individual mesophyll cell. How many protein molecules are present in a plant cell and its subcellular compartments? Which fraction of their leaf proteome do plants degrade maximally under severe drought stress? How is proteostasis controlled under these conditions? Do cells just eat anything when they are really starved or are they still picky? Are the proteins that are essential for stress resistance synthesized or rather spared from degradation? Which proteins contribute to the free amino acid pool and what happens to the amino acids released during proteolysis?

## Results

### Quantitative composition of the leaf proteome

As a starting point for investigating protein homeostasis during drought stress we focused on the proteome of control plants grown under optimal conditions to provide an impression of the status quo (Fig. 1A). Intensity-based absolute quantification (iBAQ, Schwanhäusser et al. 2011) was used for calculating the absolute content [μg protein · g^−1^ DW] of each of the 1399 different proteins detected by our shotgun-mass spectrometry approach. The complete MS dataset as well as detailed information on the calculation methods can be found in the supplemental information (Supp. Dataset S1, Supp. Fig. S1). The leaf proteome is dominated by a limited number of very high abundant proteins (Fig. 1B). RubisCO alone, which is well known for being one of the most abundant proteins on earth (Bar-On and Milo 2019), constitutes nearly one fourth of the leaf protein mass, corresponding to 26 mg · g^−1^ DW under control conditions (Supp. Dataset S1). Another fourth consists of eleven exclusively photosynthetic proteins, and in total about 80 % of the leaf protein mass can be found in the chloroplasts (Fig. 1C, top). Without taking absolute quantities into account, the distribution of the proteins detected by MS on subcellular compartments looks markedly different, with only 37 % chloroplast protein species (Fig. 1C, bottom). The protein investment of a leaf cell into different functions can be visualized on a PROTEOmap (Fig. 1D, Liebermeister et al. 2014). Under control conditions the major part of the leaf protein mass (66 %) is dedicated to photosynthesis, followed by protein metabolism (7.5 %) and amino acid metabolism (6 %).

**Fig. 1:**
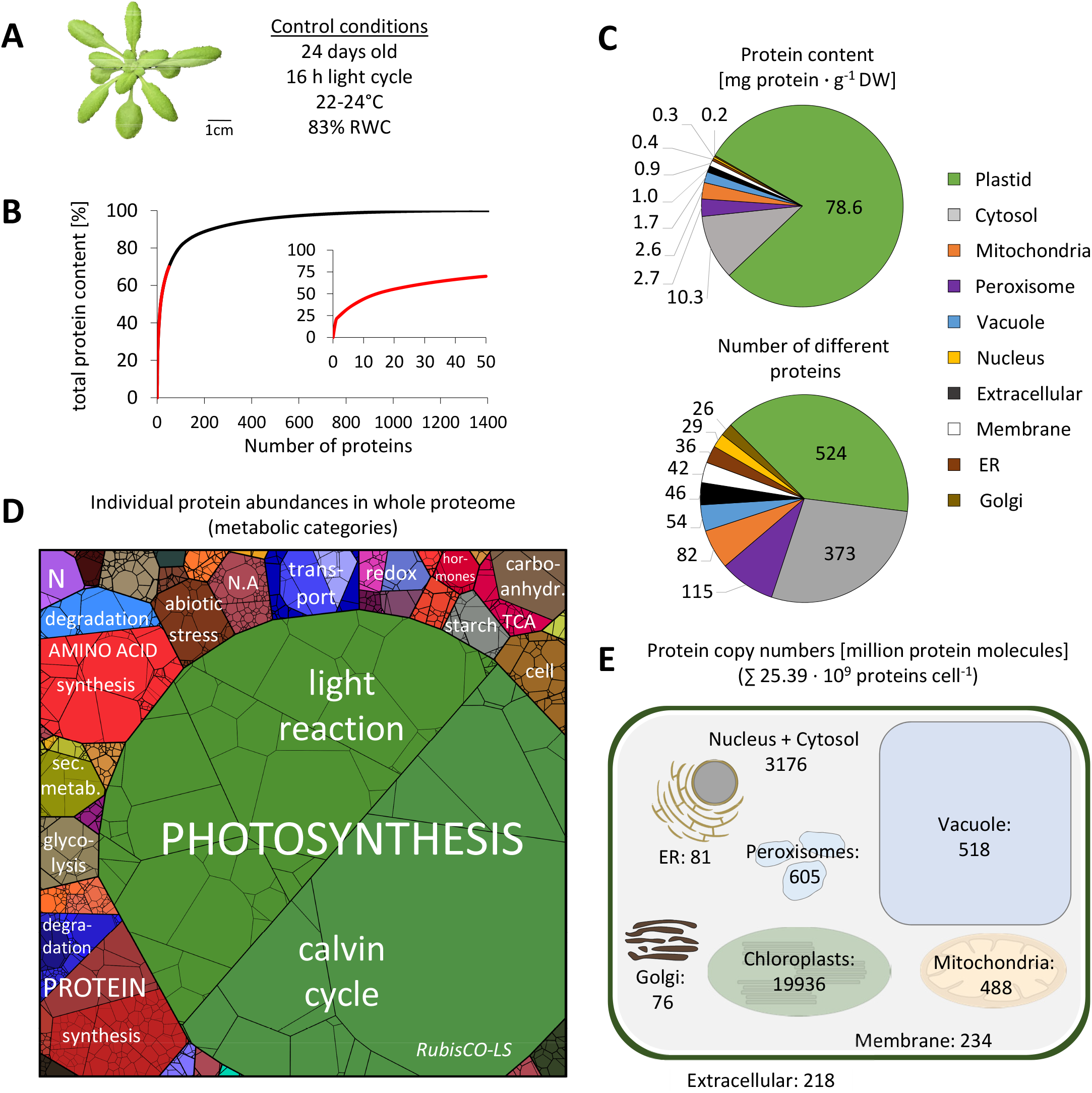
Quantitative composition of the Arabidopsis leaf proteome. Phenotype of a representative control plant used for MS analysis **B.** Fraction of total protein content contributed by each of the 1399 proteins detected by shotgun proteomics. Proteins were sorted according to their absolute content in descending order and added up. The 50 most abundant proteins (red line) are shown in the inserted graph. **C.** Distribution of the proteins detected in control samples on the different subcellular compartments according to SUBA4 (Hooper et al. 2017). Protein content (sum of all individual protein contents calculated from iBAQs) vs. number of different protein species per subcellular compartment. **D.** Proteomap illustrating the quantitative composition of the leaf proteome under control conditions. Proteins are shown as polygons whose sizes represent the mass fractions (protein abundances obtained by mass spectrometry (iBAQ), multiplied with protein molecular weight). Proteins involved in similar cellular functions according to the MapMan annotation file (version Ath_AGI_LOCUS_TAIR10_Aug2012, Thimm et al. 2004) are arranged in adjacent locations and visualized by colors. **E.** Number of protein molecules [million proteins] present in the subcellular compartments of an average Arabidopsis mesophyll cell. Copy numbers represent the sum of protein molecules present in all chloroplasts (ca. 100; Königer et al. 2008), mitochondria (300-450; Preuten et al. 2010), or peroxisomes in the cell. Copy numbers for all individual proteins detected in our MS approach are given in Supp. Dataset S1.

We used two different approaches to estimate, how many protein molecules are actually present in a plant cell based on cell number and cell size, respectively (Supp. Fig. S1, see also discussion). Both calculations consistently revealed that an average mesophyll cell in a mature Arabidopsis leaf contains about 25 billion protein molecules (Fig. 1E). 20 billion of them are localized in the chloroplasts, 3.1 billion in the cytosol, and 0.5 billion in the mitochondria. The margin of copy numbers ranges from 3.8 billion molecules of RubisCO large subunit to 2435 acetyl-CoA carboxylase 1 molecules, which is the detection limit of our MS approach. Thus, an average Arabidopsis leaf mesophyll cell contains about 400 million RubisCO hexadecamers under optimal growth conditions.

### Severe drought stress leads to massive proteolysis

We carefully established an experimental setup that mimicked physiological drought stress conditions as closely as possible and at the same time led to a highly reproducible stress phenotype (Fig. 2, a detailed description of the drought treatment is given in the methods section). Shortly, plants were grown under long-day control conditions for two weeks and watered to the same level. The dehydration process was then monitored on a daily basis and leaf samples were taken at different time points during the desiccation process from beginning to moderate and severe drought stress until recovery was no longer possible. Rosette growth gradually declined and stopped after 10 days without water (Fig. 2C). We defined this time point as stress level S1 and numbered the following days of progressive drought stress consecutively. First indications of a loss in leaf turgor became visible in some of the plants after 12 days without water (S3) and complete wilting until death occurred within the following 72 hours. These late stages of severe drought stress (S4-S7) were classified according to their leaf phenotype: Number of rolled leaves, relative water content, and potential to recover after re-watering. The leaf protein content remained stable (109 ± 13 mg · g^−1^ DW) during the first 12 days without watering (S1-S3), but then rapidly decreased by 39 % within 24 hours (S5).

**Fig. 2:**
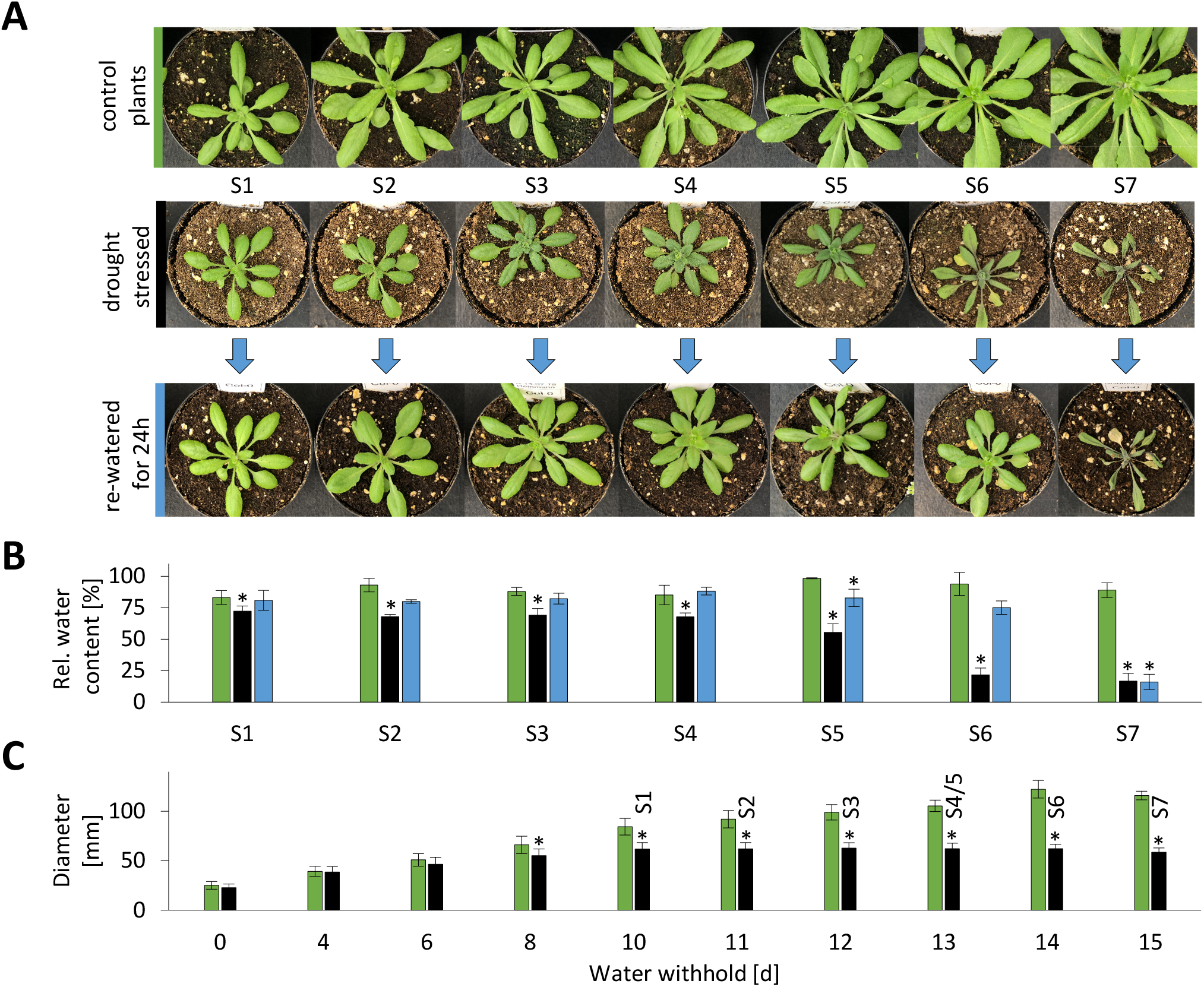
Complete setup of the progressive drought stress experiment. *Arabidopsis thaliana* plants were grown on soil under long-day conditions for 2 weeks before the start of the experiment. All pots were then brought to the same weight and the stress group was not watered for up to 15 days while the control group was kept at a constant water level. Leaf samples were taken starting after 10 days without water (stress level S1) until recovery of the plants was no longer possible (stress level S7) **A.** Phenotype of representative plants (pot diameter = 8 cm). **B.** Relative water content [%] in rosette leaves of control plants (green bars), stressed plants (black bars), and stressed plants 24h after re-watering (blue bars) at the different stress levels. **C.** Rosette diameter [mm] of control plants (green bars) and stressed plants (black bars) at 0 to 15 days after the beginning of the stress treatment. The corresponding stress levels of the plants are indicated on top of the black bars. A detailed description of the drought treatment is given in the methods section. S1-7, n=7; C1-7, n=3; R1-7, n=3. * Students t-TEST p<0.01 Starting material (stress levels) for experimental analyses:

S1 10 days after end of watering
S2 11 days after end of watering
S3 12 days after end of watering, first signs of stress (rolled/wrinkled leafs)
S4 13 days after end of watering, 4-7 rolled leaves
S5 ~ 13 days after end of watering, 8-10 rolled leaves
S6 ~ 14 days after end of watering, >10 rolled leaves; recovery of plants still possible
S7 ~ 15 days after end of watering, > 10 rolled leaves; recovery of plants not possible
C1-C7: control plants (watering continued)
R1-R7: same as S1-S7, but re-watered for 24 h

### Relative vs. absolute protein quantification during progressive drought stress

Four stress levels were selected for leaf proteome analysis by shotgun mass spectrometry (Fig 3A, Supp. Dataset S1, Supp. Fig. S2): control (relative water content (RWC) = 88 ± 5 %), S3 (moderate stress, no wilting, RWC = 69 ± 5 %), S5 (severe stress, RWC = 55 ± 7 %), S6 (maximum tolerable stress, RWC = 22 ± 5 %). Changes in the relative abundance of individual proteins were estimated via label-free quantification (LFQ) (Fig. 3B; Cox et al. 2014). This approach is suitable for identifying proteins that are induced and thus might be particularly relevant during the conditions tested. Considering the fact that plants degrade almost half of their leaf protein content during severe drought stress (Fig. 3A) it is especially important to be aware of absolute contents of the individual proteins in the leaves as well, which we calculated using iBAQ values (Fig. 3C). A combination of both data evaluation methods makes it possible to discriminate between proteins that actually increase in their absolute content even after massive proteolysis (Fig. 3B, red squares), in contrast to others that decrease less than average and as a consequence also are of higher relative abundance in the stressed plant (Fig. 3B, blue squares in the right half of the volcano plots).

**Fig. 3:**
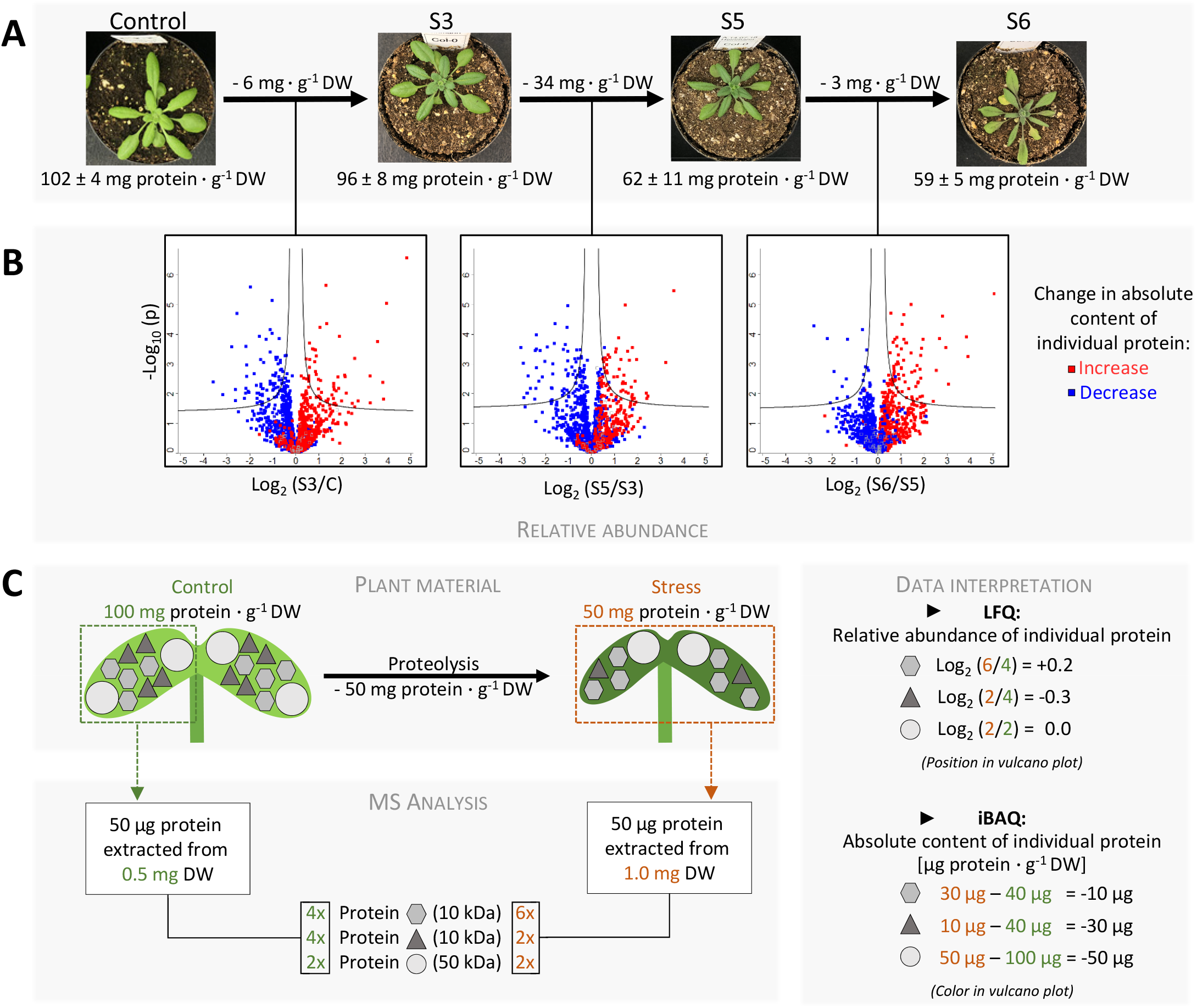
Different perspectives on the proteome of stressed plants: Relative vs. absolute changes in the leaf protein composition during progressive drought stress. **A.** Phenotype and protein content of the plants used for proteome analysis. Complete rosettes of 4 plants were harvested at the beginning of the stress treatment (parallel to S1-S3) (control), and at three defined stress stages (S3, S5, S6), respectively. **B.** Vulcano plot illustrating differences in protein abundance between the stress levels. The relative abundance of each individual protein is reflected by the position of the symbol in the plot. The curves given in solid lines represent the threshold for significance (FDR: 0.05, s0: 0.1). The symbol colors indicate changes in the absolute content of each protein (red: increase during stress, blue: decrease during stress). **C.** Schematic presentation of the two different approaches used for interpretation of the proteomics dataset. The relative abundance of each individual protein in the protein extracts used for MS analysis can be calculated on the basis of LFQ values. However, massive proteolysis during severe drought stress leads to large differences between the total protein content [mg protein · g-1 DW] of stressed vs. control plants. Therefore, in order to estimate changes in the absolute content of individual proteins during the stress treatment, we calculated the amount of each protein present in the leaf [μg protein · g-1 DW] by multiplying iBAQs with the molecular weight of the protein, calculating the mass fraction within the individual sample and multiplying it with the total protein content of the leaf. LFQ: lable-free quantification, iBAQ: intensity-based absolute quantification.

### Patterns of stress-induced proteome changes in subcellular compartments

To provide a first impression of quantitative changes in the leaf proteome during progressive drought stress we sorted all detected proteins according to their absolute content under control conditions for each compartment individually. The contents of each individual protein during progressive drought stress were then plotted in superimposing graphs (Fig. 4A). The fraction of proteins degraded in the course of the stress treatment becomes visible as green or orange area. Interestingly, there are clear differences between the compartments. A large fraction of proteins localized in chloroplasts, the cytosol, the plasma membrane, or the golgi apparatus seems to be subject to bulk degradation. In contrast, hardly any green areas are visible for mitochondrial and extracellular proteins indicating a lower degradation rate. In order to quantify this observation we calculated fold change ratios of individual protein contents in stressed vs. control plants and sorted them in ascending order for each stress level individually (Fig. 4B). Proteins with an average degradation rate are localized at 0.94 for stress level S3, at 0.61 for S5, and at 0.58 at S6, corresponding to the decrease in total protein content. In the complete dataset and also in the subsets of plastid and cytosolic proteins there is a large area with almost horizontal lines representing proteins with roughly average degradation rates. In contrast, the slopes of the mitochondrial and extracellular graphs are much steeper and only 11-12 % of the proteins show average or increased degradation rates in severely stressed plants (Fig. 4B, vertical red lines).

**Fig. 4:**
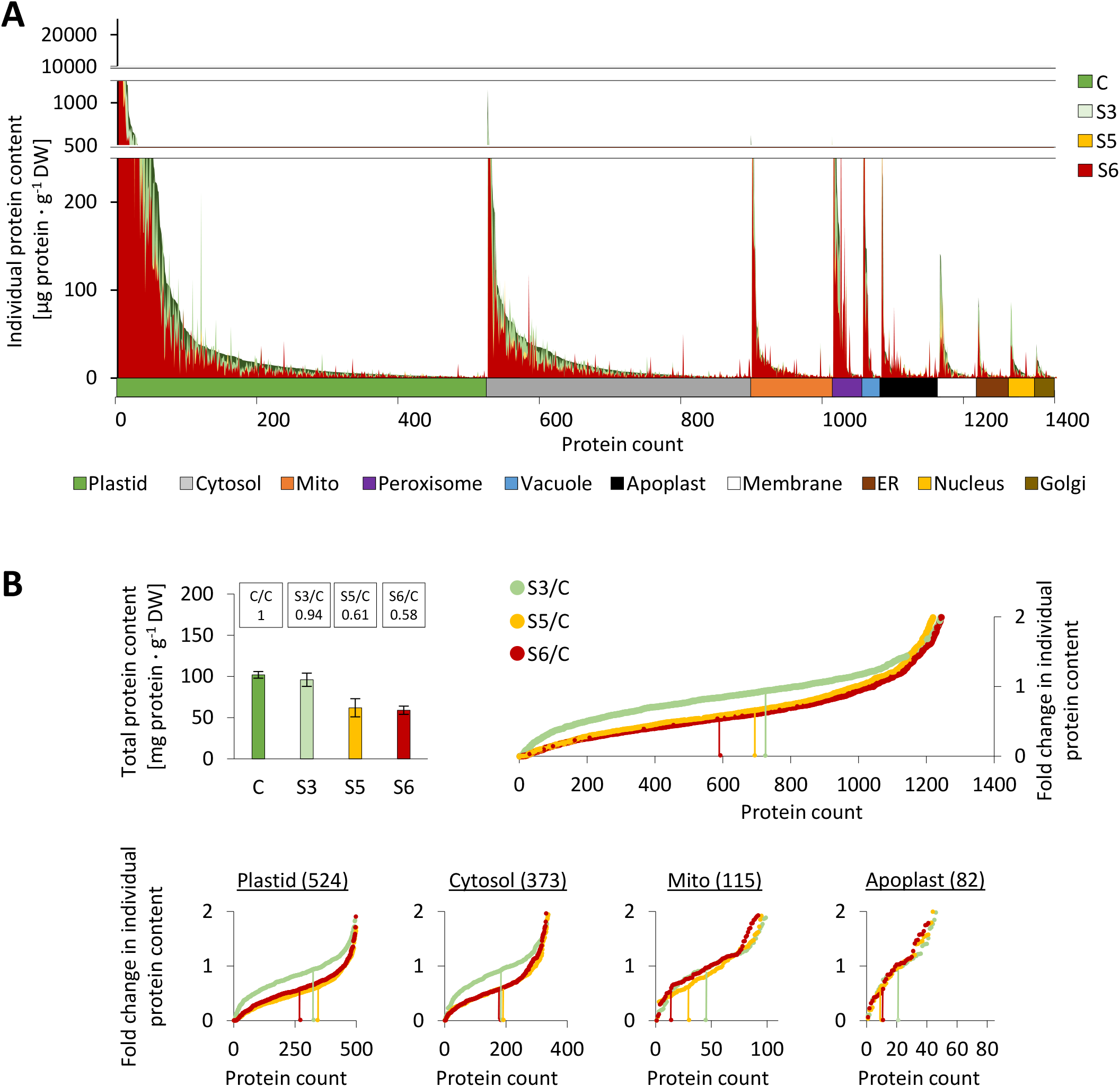
Compartment-specific patterns of stress-induced changes in individual protein abundances. **A.** Absolute contents [μg protein · g^−1^ DW] of all individual proteins detected by shotgun proteomics in descending order (under control conditions) sorted by subcellular compartments. Protein contents under control and stress conditions are shown in superimposing graphs. **B.** Fold change ratios of total leaf protein as well as individual protein contents in stressed vs. control plants. In order to visualize the fraction of proteins with average, high or low degradation rates, changes in individual protein contents were sorted in ascending order for each stress level. Vertical lines indicate proteins that correspond exactly to the decrease in total protein content, i.e. 0.94 for stress level S3 (light green), 0.61 for S5 (orange), and 0.58 at S6 (red).

### Regulation of protein abundance via synthesis and degradation

Protein abundance can be regulated at the level of synthesis and/or degradation. We used genevestigator (Hruz et al. 2008) to estimate gene expression levels during drought stress and combined this information with the relative protein abundances detected by our proteomics approach (Supp. Dataset S2). We filtered the proteomics dataset for proteins of consistently increased abundance and divided the resulting list of 332 proteins in two subgroups: Group I contained the proteins with significantly increased expression levels (88 proteins) and group II contained proteins with decreased or unaffected expression levels (244 proteins), indicating that regulation might rather be achieved at a posttranscriptional level, e.g. via decreased proteolysis (Fig. 5). In order to estimate, which metabolic pathways might preferentially be regulated by these strategies we compared the fraction of proteins attributed to a specific pathway in each regulation group to the complete MS dataset (Table 1). The proteins up-regulated via gene expression (group I) were mainly involved in protein, lipid or amino acid degradation, stress response and secondary metabolism. Energy metabolism (glycolysis and respiratory chain) and extracellular proteins required for cell wall metabolism and proteolysis were prevalent in group II and thus might be regulated by decreased degradation rates. The proteins of consistently decreased relative abundance (255 proteins) were also subdivided in those with decreased expression rates (group III, 78 proteins) and those with increased or unaffected expression rates (group IV, 177 proteins) (Fig 5, lower part). Group III (down-regulation on expression level) contains specific vacuolar proteins and enzymes catalyzing lipid or tetrapyrrole synthesis (Table 1). No particular enrichment in subcellular compartments or functional categories was detected for proteins potentially down-regulated by increased proteolysis (group IV).

**Table 1:** Estimating the enrichment of specific compartments (left) or metabolic pathways (right) in groups of proteins regulated on a transcriptional (I, III) or post-translational (II, IV) level. Numbers indicate the quotient of the fraction of proteins localized in specific compartments (left) or attributed to metabolic pathways (right) in the regulation groups (I-IV) divided by the fraction of the respective proteins in the total proteomics dataset. Group I: increased protein abundance, increased expression, group II: increased protein abundance, unaffected or decreased expression; group III: decreased protein abundance, decreased expression; group IV: decreased protein abundance, unaffected or increased expression. Only metabolic pathways with quotients ≥ 2 in at least one regulation group are shown. The complete dataset used for enrichment analysis is provided in Supp. Dataset S2.

| Compartment | I | II | III | IV | Pathway | I | II | III | IV |
| --- | --- | --- | --- | --- | --- | --- | --- | --- | --- |
| Cytosol | 1.0 | 0.8 | 0.5 | 1.1 | AA degradation | **2.6** | 0.6 | 0.6 | 1.0 |
| ER | 1.5 | 1.9 | 0.8 | 0.7 | Cell wall | 0.7 | **2.6** | 0.8 | 1.3 |
| Extracellular | **2.3** | **3.1** | 0.2 | 0.4 | Glycolysis | 1.0 | **2.2** | 0.0 | 0.0 |
| Golgi | 0.0 | 0.8 | 0.6 | 1.6 | Lipid degradation | **6.4** | 1.9 | 0.0 | 0.0 |
| Mitochondria | 1.3 | 1.8 | 0.0 | 0.3 | Lipid synthesis | 0.6 | 1.2 | **2.2** | 1.0 |
| Nucleus | **2.3** | 0.8 | 0.5 | 0.7 | mETC | 0.0 | **2.2** | 0.0 | 0.3 |
| Peroxisome | **3.2** | 0.8 | 0.0 | 1.1 | Protein degradation | **2.0** | 1.5 | 0.0 | 0.8 |
| Plasma membrane | **2.5** | 1.0 | 1.6 | 1.6 | Protein handling | 0.5 | **2.3** | 0.0 | 1.2 |
| Chloroplast | 0.3 | 0.6 | 1.7 | 1.1 | Secondary metabolism | **2.3** | 1.1 | 1.3 | 0.6 |
| Vacuole | 0.6 | 1.1 | **2.7** | 0.6 | Stress | **3.0** | 1.5 | 0.3 | 0.4 |
|  |  |  |  |  | Tetrapyrrole synthesis | 0.0 | 0.0 | **9.0** | 1.2 |

**Fig. 5:**
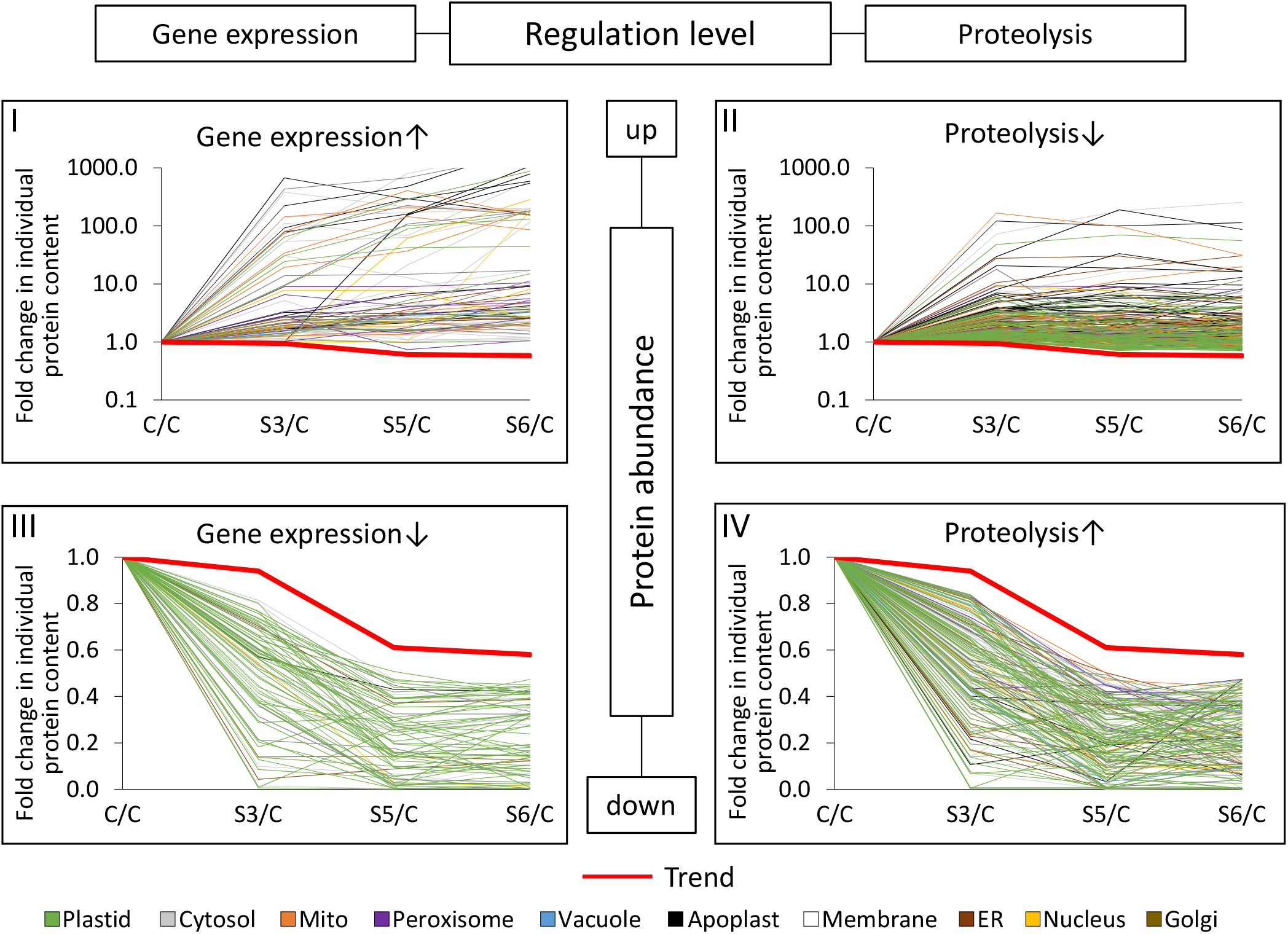
Transcriptional and post-translational regulation of protein abundances during progressive drought stress. Fold change ratios (stress/control) of individual protein contents during progressive drought stress. Red lines in the graphs (Trend) indicate the fold change in total leaf protein content at each stress level (S3/C: 0.94; S5/C: 0.61; S6/C: 0.58). Three microarray datasets available via genevestigator were used to estimate gene expression levels during drought stress (see methods section). The proteomics dataset was filtered for proteins that were of increased relative abundance according to both iBAQ-based and LFQ-based data interpretation at each stress level (more details on the filter criteria are provided in Supp. Dataset S2). These proteins of increased abundance (upper part of the figure) were divided in two groups: proteins with increased gene expression levels according to the microarray datasets (group I, top left) and proteins with unaffected or decreased gene expression levels during drought stress (group II, top right). Proteins of consistently decreased abundance (lower part of the figure) were also filtered for decreased (group III, bottom right) and increased or unaffected expression levels (group IV, bottom left). Colors indicate the subcellular localization of the individual proteins according to SUBA4 (Hooper et al. 2017). Enrichment of compartments and functional categories in the different regulation groups is listed in Table 1.

### Adaptations of the protein synthesis and degradation machineries during progressive drought stress

Under control conditions about 5.4 % of the leaf proteome detected by our MS approach is dedicated to protein synthesis (ribosomal proteins, translation initiation and elongation factors) compared to 1.4 % involved in proteolysis (proteasomes, autophagy proteins, proteases and regulatory proteins) (Fig. 6A). During progressive drought stress a majority of the proteins involved in protein synthesis (ca. 75 %) decreased more than average (Fig. 6B). In particular, the large group of ribosomal proteins (125 proteins, contributing 3.3 mg protein · g^−1^ DW under control conditions) had strikingly homogenous degradation rates (mass ratio S6/C = 0.49 ± 0.17). In contrast, the total leaf content of proteolytic enzymes remained stable (0.8-0.9 mg protein · g^−1^ DW) but changed drastically in its composition. Protease copy numbers in the cytosol, the vacuole, and in the apoplast increased progressively (Fig. 7A), and after severe stress most of the vacuolar and extracellular proteases were of significantly increased abundance compared to control conditions indicating their specific relevance for drought response (Fig. 7B). In order to estimate the mean workload of the proteolytic system in the individual subcellular compartments, we calculated the number of proteases per 1000 protein molecules (Fig. 7C). The relative abundance of proteases per substrate was at least ten fold higher in the apoplast than in any other compartment even under control conditions and further increased already during moderate stress (S3). Vacuolar proteases strongly accumulated during severe stress and also in the cytosol plus nucleus the relative capacity of proteases approximately doubled, although only a specific subset of proteolytic enzymes was significantly increased. Due to their high abundance chloroplasts contained the major fraction of cellular proteases in the leaves of non-stressed plants (Fig. 7A). However, proteases constituted less than 0.5 % of all plastid proteins (compared to 9-13 % in the apoplast) and decreased during stress to a similar extent as the majority of chloroplast proteins (Fig. 7B).

**Fig. 6:**
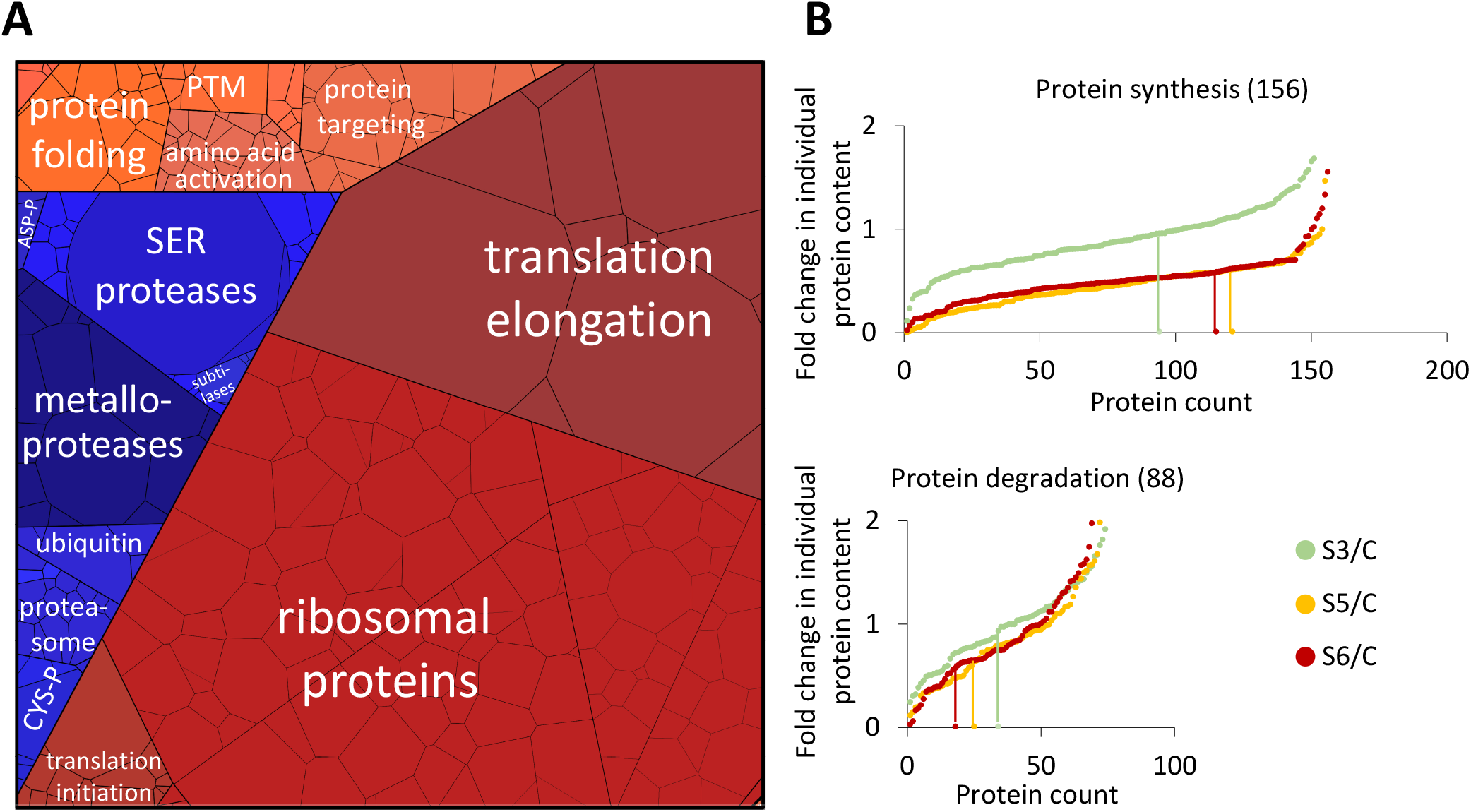
Abundance of the proteostasis apparatus during drought stress. **A.** Proteomap illustrating the quantitative composition of the proteostasis apparatus under control conditions. Proteins are shown as polygons whose sizes represent the mass fractions (protein abundances obtained by mass spectrometry (iBAQ), multiplied by protein molecular weight). Proteins involved in similar cellular functions according to the MapMan annotation file (version Ath_AGI_LOCUS_TAIR10_Aug2012) are arranged in adjacent locations and visualized by colors. The total protein fraction represented in the Proteomap is 6.9 mg · g-1 DW corresponding to 6.7 % of the leaf proteome. **B.** Fold change ratios of the individual contents of proteins involved in protein synthesis (top) or proteolysis (bottom) in stressed vs. control plants. In order to visualize the fraction of proteins with average, high or low degradation rates, changes in individual protein contents were sorted in ascending order for each stress level. Vertical lines indicate proteins that correspond exactly to the decrease in total protein content, i.e. 0.94 for stress level S3 (light green), 0.61 for S5 (orange), and 0.58 at S6 (red).

**Fig. 7:**
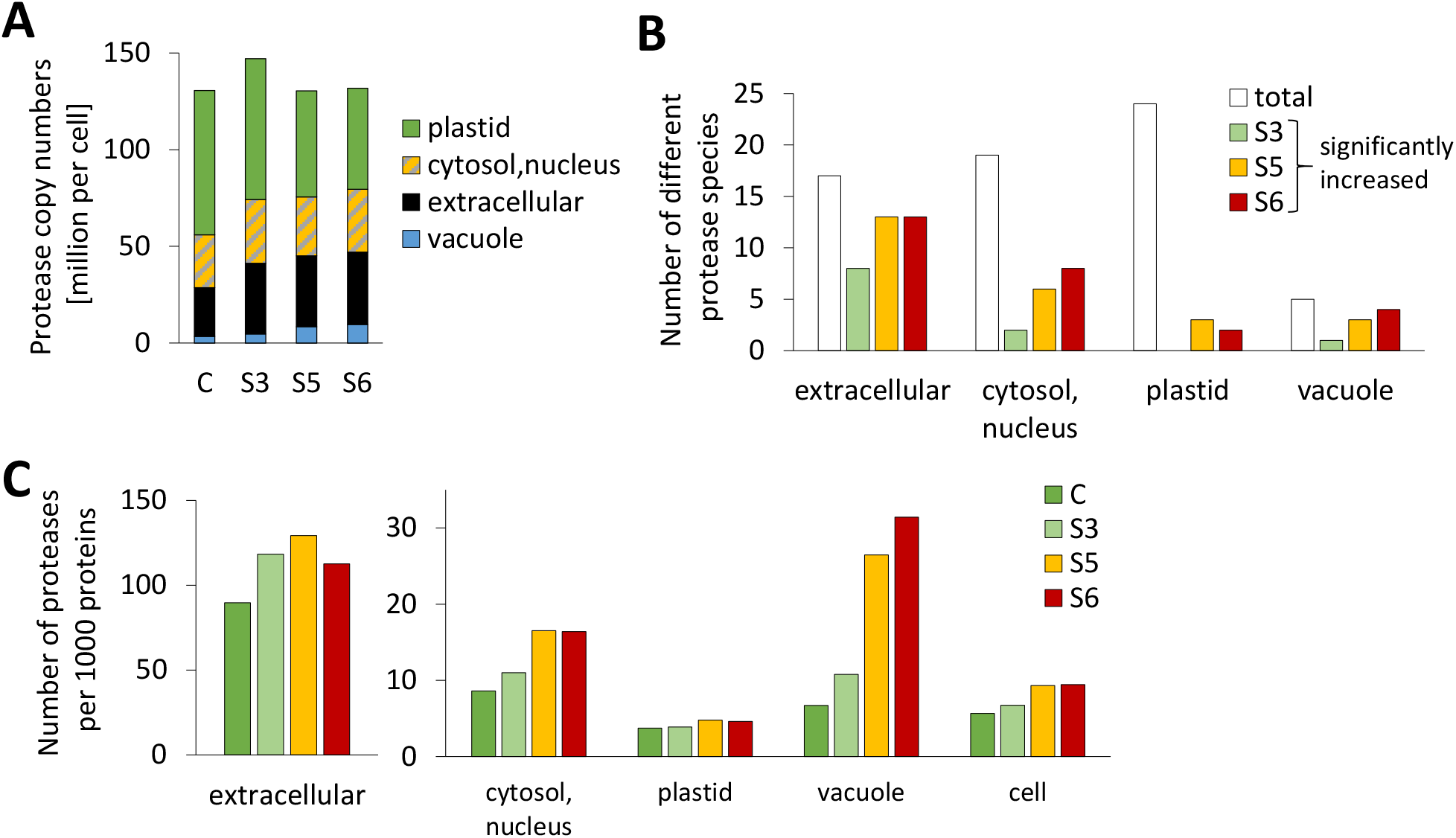
Adaptation of the proteolytic apparatus during progressive drought stress. **A.** Total number and subcellular distribution of protease molecules in an average leaf mesophyll cell under control conditions and during stress. The proteomics dataset (Supp. Dataset S1) was filtered for the MapMan category “protein.degradation” and protein copy numbers of all enzymes with proteolytic activity (without regulatory proteins and inhibitors) were added up for each subcellular compartment individually. **B.** Significant increase in the abundance of individual protease species during drought stress in the different subcellular compartments. White bars indicate the total number of different proteases detected and colored bars illustrate how many of them were significantly increased based on LFQ values at the respective stress level. **C.** Copy numbers of protease molecules per 1000 proteins in the subcellular compartments of an average mesophyll cell under control conditions and during stress. The protease copy numbers shown in A. were divided by the total number of protein molecules in the respective subcellular compartment (Fig. 1E) and multiplied by 1000.

### Dynamics in free and protein-bound amino acid pools

Massive proteolysis during severe drought stress inevitably leads to liberation of large amounts of amino acids. We thus changed perspective and focused on the further fate of the degraded part of the proteome and its effect on free amino acid homeostasis. For each individual protein we calculated the difference in absolute content in control vs. stressed plants (Fig. 8A top, Supp. Dataset S1). It immediately becomes obvious that the amino acids added to the free pool are quantitatively derived from a limited number of very high abundant proteins. Degradation of about 200 million RubisCO hexadecamers per cell alone accounts for 28 % of the total amino acid release during stress. The profiles of free amino acids in the leaves of control and stressed plants were quantified by HPLC (Supp. Dataset S3). In addition, we calculated the total amount of each individual amino acid bound in proteins on the basis of the leaf protein content and the quantitative composition of the proteome. The pool sizes and compositions of the free and protein bound amino acid pools can be visualized using a modified version of PROTEOmaps (Fig. 8B, orange: free pool, blue: protein-bound pool). Under control conditions, the Arabidopsis leaves contained 1.05 mmol · g^−1^ DW amino acids of which 0.93 mmol · g^−1^ DW were bound in proteins. Drought stress led to a decrease of the total amino content by 28 %. Also, the ratio between free and protein bound amino acids shifted from 0.13 to 0.39 due to massive proteolysis. The amino acid composition of the proteome did not change considerable during stress. The molar share of the 20 proteinogenic amino acid was in the range of 1.3 % (Cys) to 9.0 % (Ala). In contrast, the free amino acid pool strongly reacted to drought stress and also the concentrations of high and low abundant amino acids differed up to 460-fold (Fig. 8B, Supp. Dataset S4). Under control conditions, the free amino acid pool was dominated by Glu, Gln and Asp. Water deficiency led to progressive accumulation of Pro, which in the leaves of severely stressed plants represented 59 % of the free and 17 % of the total amino acid pool. In order to estimate the role of proteolysis in amino acid homeostasis we calculated the theoretical composition of the free amino acid pool that would result from partial degradation of the proteome (as detected by our proteomics approach) without any metabolic conversion of the amino acids produced (Fig. 8A, grey bars). With the clear exception of Pro the free amino acid contents actually detected in severely stressed leaves (Fig. 8A, red bars) were several fold lower than the calculated ones indicating their degradation or conversion to other metabolites. Enzymes involved in the degradation of branched-chain amino acids, Cys, Lys, and Arg were indeed increased by drought stress, as were Pro and GABA metabolism (Supp. Dataset S1, Supp. Fig. S3).

**Fig. 8:**
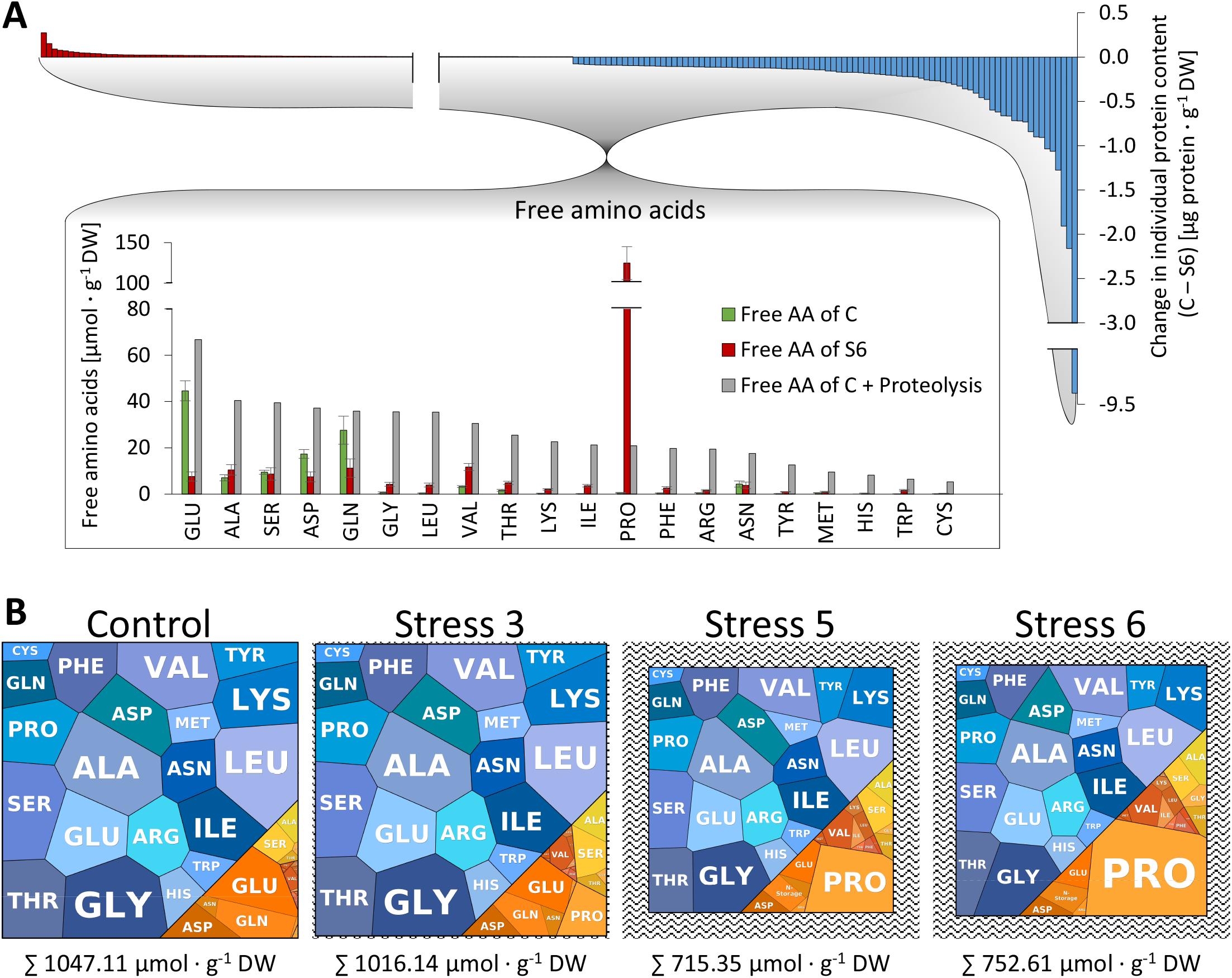
Interconnection of amino acid pools during progressive drought stress. **A.** Effect of proteolysis on free amino acid homeostasis. The quantitative composition of the degraded fraction of the proteome (blue bars) was used to calculate the theoretical composition of the free amino acid pool (grey bars) that would result from massive proteolysis during drought stress (control vs. maximum tolerable stress) without any metabolic conversion of the amino acids produced. **B.** “AMINOmaps” illustrating pool sizes and compositions of the free (orange colors) and protein bound (blue colors) amino acid pools during progressive drought stress. Amino acids are shown as polygons whose sizes represent the molar fractions. Free amino acid contents were quantified by HPLC, and quantitative amino acid composition of the proteome was calculated on the basis of molar composition of the proteome (see Supp. Dataset S1) as detailed in the methods section.

## Discussion

### Estimating protein copy numbers in a plant cell

Common sense indicates that cells require an adequate set of proteins to function properly. However, we were not able to deduce a comprehensive picture of what this protein infrastructure of a plant cell might look like from the literature. Thus we calculated the average protein copy number in a plant cell based on published information about the size and number of cells in an average Arabidopsis leaf (copy numbers are summarized in Table 2). We selected mesophyll cells as the representative leaf cell, since they are photosynthetically active and constitute the major part of the leaf volume. Total protein copy numbers have already been reported for yeast cells and different animal cell lines. A haploid cell of *Saccharomyces cerevisiae* has a volume of 42 μm^3^ (Jorgensen et al. 2002) and contains about 42 million proteins (Ho et al. 2017) whereas for human cells with a volume of about 4200 μm^3^ 3 billion protein molecules have been calculated (Kulak et al. 2014). Thus, yeast and human cells contain 1.0 and 0.7 million proteins per μm^3^, respectively. An average mature leaf cell has a volume of approximately 150.000 μm^3^ (Supp. Fig. S1). Assuming an average protein abundance of 0.85 · 10^6^ molecules per μm^3^ and subtracting the volume of the central vacuole that typically covers about 80 % of a plant cell we postulate that a leaf mesophyll cell contains about 25 billion proteins (Table 2; Supp. Fig. S1). An alternative, completely independent way to calculate protein copy numbers is based on an average number of 300.000 mesophyll cells (Wuyts et al. 2010) in a mature rosette leaf of 5 mg DW with a protein content of 102 mg · g^−1^ DW. 1.7 ng protein per cell would add up to 20.5 billion protein molecules with an average molecular weight of 50 kDa. Quantitative proteomics irrespective of its intrinsic limitations, which will be discussed in the next paragraph, makes it possible to deduce a more precise estimate of 25 billion proteins per cell and in addition it provides information about the copy numbers of individual proteins (Table 2; Supp. Dataset S1).

**Table 2:** Total number of protein molecules in an average Arabidopsis leaf mesophyll cell and its subcellular compartments under control conditions (C) and during progressive drought stress (S3, moderate stress; S5, severe stress; S6, maximum tolerable stress). For each compartment, the copy number of the most abundant protein is listed individually. All numbers are based on estimations as discussed in the text (see also Supp. Fig. S1).

|  | Protein numbers (x10<sup>6</sup>) |  |  |  |
| --- | --- | --- | --- | --- |
|  | C | S3 | S5 | S6 |
| **No. of proteins in a mesophyll cell** | **25395** | **23976** | **15069** | **15116** |
| **No. of proteins in all chloroplasts of a mesophyll cell (~100)** | **19918** | **18724** | **11457** | **11302** |
| No. of RubisCO LS (AtCg00490) per cell | 3816 | 4060 | 2424 | 2061 |
| No. of proteins in an individual chloroplast | 199 | 187 | 115 | 113 |
| No. of RubisCO LS (AtCg00490) per chloroplast | 2.0 | 1.9 | 1.2 | 1.1 |
| **No. of proteins in all mitochondria in a cell (~400)** | **488** | **473** | **380** | **401** |
| No. of serine hydroxymethyltransferase 1 (At4g37930) per cell | 65 | 50 | 31 | 29 |
| No. of proteins in an individual mitochondrion | 1.2 | 1.2 | 0.9 | 1.0 |
| No. of serine hydroxymethyltransferase 1 (At4g37930) per mito. | 0.16 | 0.13 | 0.08 | 0.07 |
| **No. of proteins in the cytosol per cell** | **2741** | **2540** | **1581** | **1672** |
| No. of GTP binding EF Tu (At5g60390) per cell | 139 | 129 | 74 | 83 |
| **No. of proteins in the vacuole** | **518** | **437** | **321** | **306** |
| No. of tonoplast intrinsic protein 2 (At3g26520) per cell | 192 | 146 | 110 | 89 |
| **No. of proteins in the extracellular space per cell** | **281** | **309** | **283** | **333** |
| No. of germin-like protein 1 (At1g72610) | 106 | 60 | 46 | 45 |
| **No. of proteins per nucleus** | **158** | **203** | **92** | **129** |
| No. of ubiquitin 5 (At3g62250) per nucleus | 37 | 49 | 21 | 21 |
| **No. of proteins in all peroxisomes of a mesophyll cell** | **605** | **641** | **491** | **525** |
| No. of Aldolase-type TIM barrel protein (At3g14415) per cell | 82 | 68 | 32 | 31 |
| **No. of proteins in the endoplasmic reticulum per cell** | **73** | **69** | **57** | **57** |
| No. of ADP-ribosylation factor 1 (At1g70490) per cell | 20 | 14 | 10 | 7.1 |
| **No. of proteins in all golgi apparatuses of a mesophyll cell** | **34** | **35** | **23** | **24** |
| No. of RGP2; UDP-arabinose mutase (At5g15650) per cell | 5.0 | 5.4 | 5.5 | 6.1 |
| **No. of proteins in the plasma membrane per cell** | **180** | **177** | **128** | **102** |
| No. of Plasma membrane intrinsic protein 2A (At3g53420) | 38 | 34 | 20 | 14 |

A major function of leaf mesophyll cells is photosynthesis and this is reflected by the large fraction of proteins (20 billion) localized in the about 100 chloroplast present in each cell corresponding to about 200 million proteins per chloroplast, and again the largest fraction of these are included in 4 million RubisCO hexadecamers (Königer et al. 2008). Interestingly, the protein copy number we calculated for the cytosol of a plant cell (3.1 million) matches almost exactly the total number of proteins reported for animal cells (Kulak et al. 2014). According to our estimation a mesophyll cell contains about 488 million mitochondrial proteins (Table 2). Assuming that between 300 and 450 mitochondria are present in a plant cell depending on the leaf age (Preuten et al. 2010), a single mitochondrion would harbor 1.1 to 1.6 million protein molecules, which is in perfect agreement with previous results (Fuchs et al. 2020).

### Strengths and limitations of the proteomics approach and its different evaluation strategies

For statistical analysis to identify significant differences between the stress levels we used LFQ, an algorithm optimized for accurate comparisons between different samples including multiple levels of normalization (Cox et al. 2014). This approach e.g. helps to identify a set of extracellular proteases that might be particularly relevant during drought stress response or to estimate the regulation of amino acid catabolic pathways (Supp. Fig. S3). However, LFQ based data interpretation is not suitable for comparing the abundance of different protein species. In contrast, a quantitative perspective on the leaf proteome based on iBAQs makes it possible to calculate mass fractions, molarities, and even copy numbers of individual proteins but it lacks statistics. Both evaluations are limited by the intrinsic shortcomings of shotgun proteomics, which cannot detect very low abundant proteins and tends to underestimate membrane proteins since the biochemical properties of their peptides such as high hydrophobicity are unfavorable for ionization and detection (Schwanhäusser et al. 2011; Fabre et al. 2014; Krey et al. 2014). Every proteomics dataset therefore has to be regarded as a representative fraction of the complete picture.

### Proteostasis under challenging conditions - individual strategies for subcellular compartments and metabolic pathways

#### How to focus on the relevant pathways during severe drought stress

Combined information about protein abundance and expression level illustrates the general strategies employed by the leaf cells to adjust their protein setup to the challenges posed by insufficient water supply. Specific stress related proteins and those involved in secondary metabolism are induced at the expression level. Similarly, cells increase the abundance of pathways that are barely used under control conditions but important to make alternative energy sources accessible such as protein, amino acid, and lipid catabolism by *de novo* synthesis. In contrast, the basic mitochondrial functions fulfilled by TCA cycle and respiratory chain are not required to be more active during stress than under control conditions, they just change their initial substrate from carbohydrates to amino acids and lipids. Therefore, it makes perfect sense that these pathways are preserved from degradation rather than up-regulated at the transcriptional level. Protection from degradation might be achieved by selective autophagy of specific organelles. During developmental senescence autophagic vesicles have been shown to preferentially contain RubisCO, entire chloroplasts, and also ribosomes whereas mitochondrial integrity and function is preserved until very late stages (Chrobok et al. 2016; Marshall and Vierstra 2018). Our results are in good agreement with this finding since we observed stronger than average decrease rates in plastid and ribosomal proteins during progressive drought stress but very little effect on mitochondrial proteins. Ribosomes are among the most stable proteins under control conditions (Li et al. 2017). However, they tie up a significant fraction of the cellular resources since they account for a majority of the cell’s RNA and also about 3 % of the protein mass. Thus, the turnover of ribosomes in eukaryotes is activated by nutritional stress such as carbon, nitrogen, or phosphate deficiency (Floyd et al. 2016). Conveniently, this measure also serves the purpose to down-regulate protein synthesis rates during stress. Apart from selective autophagy the stability of individual proteins can be regulated via ubiquitinylation and is also affected by other post-translational modifications, substrate or cofactor binding leading to faster degradation of the less busy enzymes (Nelson and Millar 2015).

#### Proteolytic systems and their contribution to stress induced protein turnover

Autophagy and proteasomes are considered to be the two major proteolytic systems in a cell. However, due to the sheer abundance of chloroplasts, the plastid proteases according to our evaluation represent the major share of proteolytic enzymes in a leaf cell under control conditions. Thus, they would be suitable for contributing considerably to the regular turnover of chloroplast proteins. Since amino acid synthesis is also localized mainly in these organelles they are perfectly equipped for exporting also the amino acids resulting from proteolysis (Pottosin and Shabala 2016). However, the frequency of proteases per total number of proteins is comparatively low in chloroplasts and in contrast to other subcellular compartments does not increase during stress. Bulk degradation of chloroplast proteins during severe dehydration therefore requires additional capacities outside the chloroplast and these can be provided by the lytic vacuoles that strongly increase their protease content and are able to hydrolyze proteins delivered by autophagic vesicles (Michaeli and Galili 2014; Marshall and Vierstra 2018).

In contrast to plastids the extracellular space is extremely rich in protease molecules per total proteins. Apart from maintaining the cell wall, major functions of the apoplast are signaling and defense against pathogens, which both involve proteolysis. Extracellular plant proteases hydrolyze proteins of invading pathogens to inactivate them and also to release signal peptides triggering immune reactions (Balakireva and Zamyatnin 2018). Plant peptide hormones are usually produced as pre-pro-protein and need to be activated by proteolytic cleavage (Stührwohldt and Schaller 2019). This function has been shown to be particularly relevant for drought resistance. Extracellular subtilisin-like proteases are involved in the regulation of stomatal density and distribution in response to environmental stimuli (Berger and Altmann 2000; Engineer et al. 2014). In addition, the subtilase SASP degrades and thus inactivates OST1, a kinase activated by abscisic acid (ABA), and therefore acts as a negative regulator in ABA signaling (Wang et al. 2018). Our dataset shows a strong induction of SASP during drought stress and identifies 14 additional extracellular proteases that are significantly increased and thus might be relevant for stress resistance. The apoplast proteome is remarkably stable even during severe dehydration. This finding might indicate a specific relevance of extracellular proteins during drought stress, which is clearly the case for proteases. An alternative explanation could be that apoplast proteins simply evade the intracellular bulk degradation systems autophagy and proteasome due to their remote localization.

### Amino acid homeostasis under challenging conditions – massive adjustments to the free pool provide osmolytes and ATP

The free pool represents only about 11 % of all cellular amino acids under control conditions but strongly gains impact in the course of the drought stress response. Also, despite massive proteolysis the relative composition of the proteome looks roughly similar before and after stress (Supp. Fig. S4), whereas changes on the metabolite level are rapid and drastic (Fig. 8B). Taken together these observations illustrate that homeostasis has a different meaning with regard to free amino acids and proteins. Proline is a well-known compatible osmolyte in plants and also in some euryhaline animals (Szabados and Savouré 2010; Wiesenthal et al. 2019). Free proline accumulated 219 fold and even its total amount (free plus bound in proteins) increased from 48 to 153 μmol ⋅ g^−1^ DW during progressive drought stress indicating extensive *de novo* synthesis (Supp. Dataset S4, Supp. Fig. S5). In contrast, the total contents of all 19 proteinogenic amino acids except Pro clearly decreased during the stress phase indicating that they are most likely not synthesized during stress, but accumulate in the free pool as a consequence of proteolysis (Supp. Fig. S5). An exception might be those amino acids that serve as precursors for secondary metabolites such as the aromatic amino acids (Tzin and Galili 2010). The sum of all amino acids dropped by 29 % during stress most likely due to their use as alternative respiratory substrates and precursors for secondary metabolites (Araujo et al. 2011; Hildebrandt 2018). In order to develop an idea about how long plants would be able to keep up their regular mitochondrial respiration rate when using exclusively the amino acids released by protein degradation as substrates we calculated the total number of electrons that would be transferred to oxygen via the mitochondrial respiratory chain during complete oxidation of the specific set of amino acids released during drought stress (Supp. Dataset S4, Hildebrandt et al. 2015). This oxidation process would lead to a total oxygen consumption of 1062 μmol O_2_g^−1^ DW and thus, on the basis of a mean leaf respiration rate of 3.4 nmol O_2_ · g^−1^ fresh weight · s^−1^ (O’Leary et al. 2017) could fully sustain leaf energy metabolism for about seven hours. However, leaf respiration rates tend to decrease during dehydration (Pinheiro and Shaves 2011), so that amino acid catabolism in addition to some residual photosynthetic activity and the oxidation of lipids and chlorophyll can be anticipated to make a significant contribution to the ATP supply of drought stressed plants.

## Materials and Methods

### Plant growth and drought stress treatment

#### Arabidopsis thaliana

Columbia-0 plants were grown for two weeks in pots (200 cm^3^) in a phytochamber (22 - 24 °C, 16 h light, 8 h darkness, 110 μmol s^−1^ m^−2^ light). The stress treatment started with soaking the substrate (Steckmedium, Klasmann-Deilmann GmbH) with tap water to a distinct weight (150 g). A uniform desiccation process was achieved by monitoring pot weights and reorganizing the positions of the pots in the chamber every other day. After 10 days without watering leaf material (complete rosettes) was harvested on a daily basis (Fig. 2). During late stages of severe drought stress (S4-S7) plants were additionally sub-classified according to their leaf phenotype (S4: 4-7 rolled leaves S5: 8-10 rolled leaves, S6: > 10 rolled leaves). For each stress level, seven stressed plants and three controls were harvested. In addition, three stressed plants were re-watered to test their viability and harvested after 24h.

### Determination of relative water content (RWC)

The method used is based on Smart and Bingham 1974. The weight of a leaf was measured immediately after harvest (fresh weight, FW), after overnight incubation in distilled water (turgor weight, TW), and after overnight drying at 37 °C (dry weight, DW).

RWC was calculated according to the following formular: 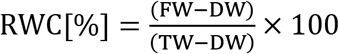

### Extraction and quantification of total protein

5 mg lyophilized plant rosette powder was dissolved in 700 μl methanol (100 %) and incubated for 20 min shaking at 80 °C. After centrifugation (10 min, 4 °C, 18.800 xg) the pellet was washed twice in 1 ml ethanol (70 %) and resuspended in 400 μl NaOH (0.1 M). The solution was incubated for 1h shaking at 95 °C and centrifuged again. The protein content of the supernatant was quantified using Ready-to-use Coomassie Blue G-250 Protein Assay Reagent (ThermoFisher) and Albumin Standard 23209 (ThermoFisher).

### Quantification of free amino acids by HPLC

Free amino acids were extracted as described in Batista et al. (2019). The pre-column derivatization with o-phthaldialdehyde (OPA) and fluorenylmethoxycarbonyl (FMOC) was based on the application note “Automated amino acids analysis using an Agilent Poroshell HPH-C18 Column” by Agilent. The samples were injected onto a 100 mm x 3 mm InfinityLab Poroshell HPH-C18 column (2.7 μm) using an Ultimate 3000 HPLC system (ThermoFisher). HPLC settings were set as described in Batista et al. 2019. Cysteine was quantified after derivatization with the fluorescent dye monobromobimane using the same HPLC system (Fahey et al 1980; Newton et al. 1981). 5 mg lyophilized plant powder was mixed with 10 μl bromobimane (46 mM in acetonitrile), 100 μl acetonitrile, and 200 μl buffer (160 mM HEPES, 16 mM EDTA, pH 8.0) and incubated on a shaker for 30 min in darkness before adding 100 μL methanesulfonic acid (65 mM). Samples were separated on a LiChrospher 60 RP-select Hibar RT 5 μm column (Merck) at 18 °C using a gradient of two solvents (0.25 % acetic acid (pH 4) and methanol). Labeled thiols were detected using a fluorescence detector 3400 RS (ThermoFisher) at 380 nm for excitation and 480 nm for emission.

### Protein extraction and label-free quantitative shotgun mass spectrometry

For protein extraction, about 5 mg of the lyophilized rosette powder was used (C, S3, S5, S6; n=4). Protein extraction, sample preparation, and LC-MS/MS were performed as previously described (Thal et al. 2018) using a Q-Exactive mass spectrometer coupled to an Ultimate 3000 UPLC (ThermoFisher).

### Protein identification by MaxQuant and data processing via Perseus software

The LC-MS/MS spectra were analyzed using MaxQuant (Version 1.5.5.1, Cox and Mann 2008) and protein identification was based on the TAIR10 database. The search parameters were set to: carbamidomethylation (C) as fixed modification, oxidation (M) and acetylation (protein N-term) as variable modifications. The specific digestion mode was set to trypsin (P) and a maximum of two missed cleavage sites was allowed. FDR at the protein and PSM level was set to 1 %. For maximum proteome coverage, the minimum number of unique peptides per protein group was 1. Unique and razor peptides were used for protein quantification. The iBAQ function of MaxQuant was enabled, “log fit” disabled. Further analysis and statistical evaluation based on LFQ and iBAQ values generated by MaxQuant were performed in Perseus (version 1.6.1.1), (Tyanova et al. 2016). The LFQ dataset was filtered to remove potential contaminations, reverse sequences or those only identified by site. Proteins were also excluded from further analysis if they were not detected in at least three of four replicates in at least one group (C, S3, S5, S6). Missing protein intensities were then considered as too low for proper quantification and replaced by very low values from a normal distribution. Finally, a list of 1399 proteins (Supp. Dataset S1) was used for all further calculations. Statistical analysis of the MS dataset was performed in Perseus using two-sample t-tests (P < 0.05).

### Calculating absolute contents of individual proteins based on iBAQ values

Raw iBAQ values generated by MaxQuant were multiplied with the molecular weight of the respective protein [kDa]. These individual weighted iBAQs were then divided by the sum of weighted iBAQs of all detected proteins for normalization and means of the four biological replicates in each sample group were calculated. The mean mass fractions were then multiplied with the total protein content of the sample [mg · g^−1^ DW] to determine the mass content of each individual protein [μg · g^−1^ DW]. The mass contents were divided by the molecular weight of the respective protein to calculate the molar protein contents [nmol · g^−1^ DW]. Protein copy numbers in an individual mesophyll cell were calculated by multiplying the molar protein contents with the mean leaf dry weight and the Avogadro constant and dividing it by the mean number of mesophyll cells per leaf. A more detailed description of the calculation methods is provided in Supp. Figure S1.

### Calculating protein bound amino acid contents based on individual protein contents

The amino acid composition of each protein was determined on the basis of its sequence. The molar content of the protein was then multiplied with the number of each of the 20 amino acids present in this protein to calculate the molar contents of the individual amino acids. The resulting molar amino acid contents were summed up for all identified proteins in a sample. The total numbers of amino acids released due to proteolysis were calculated by subtracting contents of protein bound amino acids in stressed and control plants.

### Calculating mitochondrial oxygen consumption with amino acids as alternative respiratory substrates

To estimate mitochondrial respiration in leaves that use exclusively the set of amino acids released by protein degradation during drought stress as substrates total leaf amino acid contents of stressed plants were subtracted from those of control plants. For each amino acid this difference was multiplied with the number of electrons transferred to the respiratory chain during complete oxidation (Hildebrandt et al. 2015) and divided by four to calculate the total amount of oxygen consumed (Supp. Dataset S4).

### Genevestigator datasets

The following three microarray datasets were used for estimating gene expression levels during drought stress: 1. AT-00684_1 (Ludwikow et al. 2009; long-day conditions, start: 3 weeks, samples after 5 days of dehydration in soil); 2. AT-00626_1 (Pandey et al. 2013: long-day conditions, start: 3 weeks, samples after 10 days of dehydration in soil); 3. AT-00292_1 (Perera et al. 2008: short-day conditions, start: 6 weeks, samples after 7 days of dehydration in soil)

## Supplemental Data Files

Supplemental Figure S1: Calculation of individual protein contents and copy numbers

Supplemental Figure S2: Principal component analysis of the MS dataset

Supplemental Figure S3: Drought stress-induced amino acid degradation delivers nitrogen and glutamate for the production of proline and GABA as osmolytes

Supplemental Figure S4: Changes in the quantitative composition of the leaf proteome during drought stress (Proteomaps)

Supplemental Figure S5: Sum of free and protein bound contents for all 20 proteinogenic amino acids during progressive drought stress in *Arabidopsis thaliana* rosette leaves

Supplemental Dataset S1: Complete MS dataset: LFQ and iBAQ values, relative protein abundances, mass contents [μg · g^−1^ DW], molar contents [nmol · g^−1^ DW], and copy numbers [million proteins per cell] of 1399 protein species during progressive drought stress

Supplemental Dataset S2: Combined analysis of protein abundances and expression levels to identify general strategies of leaf cells to adjust their protein setup to the challenges posed by insufficient water supply.

Supplemental Dataset S3: Free amino acid contents in *Arabidopsis thaliana* rosette leaves during progressive drought stress.

Supplemental Dataset S4: Pools of bound and free proteinogenic amino acids in *Arabidopsis thaliana* rosette leaves during progressive drought stress.

## Acknowledgements

We thank Marianne Langer and Dagmar Lewejohann for skillful technical assistance during MS sample preparation and Michael Senkler for IT support.

## Author Contributions

TMH and HPB initiated the project; TMH designed the research; BH performed most experiments; BH and PK performed the shotgun proteomics experiments; TMH and BH analyzed the data; TMH wrote the paper with support from HPB and BH.

